# Transcriptomic Changes and Biomarkers in Barrett’s Metaplasia, Dysplasia and Cancer

**DOI:** 10.64898/2026.08.25.746910

**Authors:** Thejaani P. Udumanne, Yi Jin Liew, Dana Pascovici, Tao Yang, Ka Ki Michelle Lee-Ng, Gary Gracie, Priyanthi Kumarasinghe, Duncan McLeod, Ian Brown, Michael J. Bourke, Sarah J. Lord, Jason Ross, Reginald V. Lord

## Abstract

Esophageal adenocarcinoma (EAC) has a poor five-year survival rate and one of the fastest-rising incidences of any cancer. The presence of dysplasia in Barrett’s esophagus (BE) is the main risk factor for EAC development and guides clinical management. Unfortunately, the current histopathological diagnosis of dysplasia is unreliable, with poor inter-observer agreement, highlighting the need for novel biomarkers that can improve diagnostic accuracy. Here, we performed transcriptome profiling across the full spectrum of BE-related neoplasia in 85 samples to delineate gene expression alterations in progressively worse disease stages and identify biomarkers that could complement histopathology to improve the detection of dysplasia and EAC in endoscopic biopsy specimens. Differential gene expression and pathway analyses revealed that the most extensive transcriptional changes occurred during the transition from normal squamous (NSq) to non-dysplastic BE (NDBE), consistent with metaplastic transformation. Compared to NDBE, dysplasia was characterized by enhanced cellular growth and proliferation; upregulation of immune processes and oncogenic signaling pathways were present in EAC. Using machine learning approaches, we identified a novel five-gene panel suitable for a potential RNAseq-based diagnostic test (SLC11A1, IL36A, LUCAT1, MIR215, RNU6-954P) and performed an initial validation of this signature in an additional 51 samples. We also identified several potential novel immunohistochemical markers that may warrant further evaluation, including TREM1, CXCL5, OSM, and motilin. In summary, by delineating transcriptional changes across the full disease spectrum, this study identifies several candidate biomarkers for improving current diagnostic methods for Barrett’s dysplasia and EAC.

**Significance statement:** Current histopathological assessment of Barrett’s dysplasia has limited diagnostic reproducibility. By defining transcriptional changes across the disease spectrum, this study identifies novel biomarker candidates for the detection of dysplasia and adenocarcinoma. These include a five-gene panel for a potential RNA sequencing-based diagnostic test to complement histopathology.

## Introduction

In Barrett’s esophagus (BE), the normal squamous lining of the distal esophagus is replaced by a metaplastic intestinal-like columnar epithelium containing mucous-secreting goblet cells. The development of BE is a response to long-standing gastroesophageal reflux disease (GERD) and is estimated to occur in 3% of adults in Western and 1% in non-Western countries (1). While most patients with BE have stable non-dysplastic BE (NDBE, intestinal metaplasia (IM)), management includes endoscopic surveillance because a minority progress to the stages of low-grade dysplasia (LGD), high-grade dysplasia (HGD), and esophageal adenocarcinoma (EAC).

EAC has one of the highest case-fatality ratios of all cancers (5-year survival rate approximately 20%) (2) and possibly the highest increase in incidence, with an over 6-fold incidence rise in some regions in the last four decades (3). The presence of dysplasia in BE is the major risk factor for EAC development and determines the management of this disease. Patients with NDBE have a very low annual risk of progression to EAC, estimated in a meta-analysis at 0.33% per year (4). The risk is substantially higher in patients with dysplasia, but for LGD in particular, the estimates of the risk of progression to HGD/EAC vary widely in different reports, from <1.0% to >10% per year (5,6). This likely reflects variability in the histopathological diagnosis of LGD, which is often difficult and unreliable, with studies showing poor reproducibility (7,8).

Recognition of the cancer risk associated with LGD has led to endoscopic eradication therapy (EET) being used for LGD at some centers, whereas in the past EET was reserved for HGD and superficial EAC, such as intramucosal cancer (IMC, T1a EAC) (9). The presence of dysplasia also guides surveillance endoscopy intervals for BE. Together, the clinical relevance of dysplasia and the unreliability of histopathological diagnosis support the potential value of novel biomarkers that can be added to conventional histopathology to improve diagnostic accuracy for dysplastic BE.

To date, reported diagnostic biomarkers from biopsied tissues or collected cells primarily distinguish BE and EAC from the normal squamous esophageal lining (NSq). These include the protein marker TFF3, DNA methylation markers (TFPI2, CCNA1, VIM, p16), chromosomal abnormalities (chr7 and 17, 8q24 (C-MYC), 17q12 (HER2), 20q13), and microRNAs (miRNA 192, 195, 215) (10,11).

Over the past two decades, gene expression studies of BE/EAC have evolved alongside advances in sequencing technologies, progressing from microarray-based techniques to bulk RNA sequencing to more recently single cell and spatial transcriptomics (12,13). Many of these studies have focused on the following areas: (1) comparing BE and/or EAC with NSq tissue to identify disease specific alterations (14,15), (2) comparing EAC with BE to identify changes associated with malignant transformation (16,17), (3) EAC specific studies on molecular subtyping, tumor microenvironment and prognosis (18,19), (4) disease progression and risk stratification in BE (20,21), and (5) the cellular origin of BE and EAC development (22,23). However, perhaps because of the diagnostic difficulty with LGD, fewer studies have investigated transcriptional changes across the entire disease spectrum, from NSq tissue through Barrett’s metaplasia (NDBE) and dysplasia to EAC.

Our previous whole transcriptome RNA-sequencing study (24), utilizing NSq, NDBE, LGD, and EAC samples, characterized dysregulation of long non-coding RNAs and repeat elements that may contribute to disease progression. In the present study, we expanded the number of patients and tissues used for transcriptomic profiling and included patients with HGD. We aimed to characterize stage-wise gene expression changes across the full disease spectrum and identify potential transcriptomic biomarkers that could complement histopathological methods to distinguish LGD, HGD, and EAC from NDBE.

## Materials and Methods

### Study cohort and sample selection

The study cohort included 71 patients undergoing endoscopy enrolled in the Progression of Barrett’s Oesophagus to Cancer Network (PROBE-NET) study, with fresh frozen esophageal biopsy tissue samples available. The eligibility criteria and PROBE-NET study details have been reported previously (25). Eighty-five tissue samples were included for RNAseq analysis: NSq (biopsy at 20cm from incisors)=10, NDBE=29, LGD=21, HGD=11, EAC=14. The findings were validated using a published dataset consisting of 51 samples (NSq=17, NDBE=14, LGD=8, EAC=12) from 44 patients (24) (raw expression data from Maag et al. (2017) study are available at https://www.ebi.ac.uk/ena/ under accession number PRJEB11797).

Ethics approval was obtained through the PROBE-NET study (2019/ETH03452). Written informed consent was obtained from all participants prior to recruitment and sample collection.

### Specimen collection and disease classification

Tissue samples were immediately placed in RNAlater solution (Thermo Fisher Scientific, cat. #AM7020) for stabilization. The samples were stored at 4°C for 12-24 hours to ensure tissue preservation, then transferred to −80°C for long-term storage. The central section of each specimen was formalin-fixed, paraffin-embedded and processed for Hematoxylin and Eosin (H&E) staining and subsequent histopathological assessment. The H&E section diagnoses required the agreement of at least two expert BE pathologists.

### RNA extraction, library preparation and sequencing

Total RNA was extracted and purified from homogenized tissue using RNeasy Mini Kit (Qiagen, cat. #74104) according to the manufacturer’s instructions. Prior to extraction, fresh frozen biopsy tissues were homogenized on ice in the supplied lysis buffer using a handheld motorized homogenizer with disposable pestles. DNase-I (Qiagen, cat. # 79254) treatment was performed prior to elution. RNA concentration in each sample was quantified using QuantiFluor® ONE dsDNA System (Cat. #E3310). The purity of the samples was assessed using Implen NanoPhotometer. RNA integrity was monitored on TapeStation (Agilent) using RNA Screen Tape (Integrated Sciences, cat. #5067-5576).

Total RNA isolated was sequenced at the Australian Genome Research Facility (AGRF). The libraries were prepared using Illumina Stranded Total RNA Prep with Ribo-Zero Plus kit and sequenced on NovaSeq X platform using a paired-end 2x150bp setting.

### RNA-seq data analysis

Sequencing data alignment was carried out using STAR aligner (v2.7.11b) (26), according to the parameters defined in Genomic Data Commons mRNA Analysis Pipeline (27) using the human reference genome GRCh38. Gene-level read counts were generated using STAR’s built-in quantification mode against GENCODE v36 annotations. Transcripts Per Million (TPM) counts for all samples were computed using Salmon (v1.10.1) (28). For publicly available datasets, raw FASTQ files were downloaded and processed using the same pipeline described above.

For each sample, tissue purity scores along with levels of immune cell infiltration and stromal cell presence were assessed *in-silico* using Estimation of STromal and Immune cells in MAlignant Tumors using Expression data (ESTIMATE) algorithm (29), implemented via tidyestimate R package (v1.1.1). Differential gene expression analysis was performed on gene-level counts using the DESeq2 (v1.46.0) (30), using a significant cutoff of |log2 fold change| > 1, adjusted p-value < 0.05. Pairwise comparisons of groups ordered by extent of disease progression stage were evaluated (NDBE vs. NSq, LGD vs. NDBE, HGD vs. LGD, EAC vs. HGD), together with contrasts comparing all disease stages to NSq and NDBE. As an additional analysis, the immune and stromal scores calculated using ESTIMATE were incorporated as covariates in the DESeq2 model (adjusted model).

Gene Ontology (GO, Biological Process) (31) enrichment analyses were performed using clusterProfiler (v4.14.6) (32). For pair-wise comparisons across increasing disease severity, Gene Set Enrichment Analysis (GSEA) was conducted on ranked lists of differentially expressed genes (33), whereas Over-Representation Analysis (ORA) was used for disease stage comparisons with NSq (34). Additional pathway analyses were performed using decoupleR (v2.12.0) (35) to assess Pathway RespOnsive GENes (PROGENy) model (36) and transcription factor (TF) enrichment using CollecTRI network (37).

### Biomarker selection

Candidate biomarkers were identified for protein-coding and non-coding genes using differentially expressed gene lists generated by DESeq2 for selected pairwise and group comparisons across increasing disease severity from NDBE to dysplasia to EAC. Only genes consistently identified as upregulated or downregulated in both simple and adjusted models were utilized and a |log2 fold change| > 2 and adjusted p-value < 0.05 were used as cutoffs to assess significance.

To further refine the biomarker list, expression-based filtering was applied in relevance to dysplasia and EAC as follows. Upregulated genes were required to have a median TPM > 5, while downregulated genes were required to have a median TPM = 0 in the worse disease stages in the respective pairwise comparison (i.e. group of interest). In addition, genes were filtered to only retain the ones that demonstrated a ratio of average normalized counts greater than 5 between the group of interest and the reference group.

The genes meeting all criteria were shortlisted for validation and were first assessed using the STRING database (38) to examine protein-protein interactions (PPI) to ensure biological relevance. For the top 5 clusters identified in STRING network analysis, the median area under the curve (AUC) was calculated for both discovery and validation datasets using pROC (v1.19.0.1) (39).

### Machine learning based ranking of biomarkers

Variable selection was performed to rank biomarkers that can distinguish between controls (NSq/NDBE) and cases (HGD/EAC), with LGD alternatively included in each group, based on the scores generated using LASSO regression (40) (glmnet v5.0) and XGBoost (41) (xgboost v3.2.1.1). The selection was repeated 100 times on random subsamples of data, i.e. five-sixth of data with replacement. Top 20 features ranked by importance from each method were extracted and combined to generate an overall ranking. The diagnostic performance of the selected biomarkers was assessed by calculating the AUC in the discovery and validation cohorts separately.

### Panel optimization for RNAseq based test development

Starting with the top ranked marker, features were added sequentially, and performance was evaluated at each step using a logistic regression model with leave-one-out cross validation (LOOCV). The discriminative ability of the selected markers was quantified using AUC and markers were added iteratively, selecting at each point the feature maximizing the resulting AUC, until the improvements became marginal (less than 0.005). The process continued until no further improvement was observed. The final panel was evaluated in the validation cohort.

### Immunohistochemistry (IHC) validation

Following machine learning-based ranking and AUC calculations, nine targets were selected for validation using immunohistochemistry (IHC), alongside three established clinical biomarkers. Three commercially available tissue microarrays (TMAs) containing formalin-fixed, paraffin-embedded (FFPE) 56 EAC (including gastroesophageal junction adenocarcinoma) and 23 NSq esophageal tissue were utilized for IHC validation experiments (Tissue Array, cat. #ES8011c, #ES781 and #GI501).

IHC staining was performed using the Ventana BenchMark Ultra automated staining platform (Roche Diagnostics) using manufacturer’s instructions. Briefly, the TMA slides were baked at 60°C, deparaffinized, and subjected to heat-induced epitope retrieval, with specific conditions optimized for each antibody. Details of the IHC methods for each antibody are summarized in the Supplementary Table S1. For all antibodies, except for Ki67, TMA staining was recorded with respect to staining intensity (range of 0-3) and percentage of positive cells (0-100%), which were then used to calculate H-score (range of 0-300). For Ki67, only the percentage of positive cells was recorded. Where duplicate or triplicate cores were available, H-scores or the percentage of positive cells were averaged for each case. Statistical significance was evaluated using Wilcoxon rank sum test with continuity correction.

### Data and code availability

Raw FASTQ files generated in this study have been deposited in the European Nucleotide Archive (ENA, https://www.ebi.ac.uk/ena) and will be released publicly upon publication. Processed data supporting the findings are available from the corresponding author upon request. The bioinformatics pipeline and custom scripts used for data processing and analysis are available at GitHub repository: https://github.com/tudumanne/BulkRNAseq_Barretts.

## Results

### Study cohort and RNA sequencing overview

The selected patient samples were from 62 (87%) males and 9 (13%) females, with a median age of 66 years (range from 29 to 85 years). The average number of reads generated for 85 sequenced samples was ∼51 million paired reads, with all samples achieving >80% uniquely mapped reads. Detailed cohort characteristics and RNA sequencing run summaries are provided in Supplementary Table S2.

### Tissue composition minimally influenced differential expression analyses

We performed *in-silico* tissue purity estimations using ESTIMATE (29), which first calculates immune and stromal scores for each sample by applying single sample gene enrichment analysis (ssGSEA) using predefined gene signatures. These two scores are then combined and transformed into tumor purity values ranging from 0 to 1 using a nonlinear regression model (Supplementary Figure S1A-C). The underlying assumption is that higher stromal and immune infiltration corresponds to lower tumor purity and vice versa (29). Tissue composition of biopsy samples can vary due to biological heterogeneity, microenvironment, technical or sampling bias. Therefore, while NSq, NDBE, and dysplasia samples do not contain tumor cells, we applied this method across all samples to approximately assess the extent of stromal and immune cell presence, and the overall purity.

In terms of stromal and immune scores, a steadily increasing trend across disease severity was observed for the stromal score while the immune cell component decreased from NSq to NDBE/LGD, before increasing in HGD and EAC (Supplementary Table S2, Supplementary Figure S1A-B). ESTIMATE analysis resulted in purity scores between 0.59-0.96, with relatively lower purity observed in EAC samples compared to NSq and NDBE/dysplasia samples (median NSq=0.89, NDBE=0.91, LGD=0.89, HGD=0.90, EAC=0.80, Supplementary Figure S1C). This is consistent with the higher stromal and immune scores observed in our study and with the well-described cellular composition of EAC consisting of immune and stromal cell populations, alongside malignant epithelial cells (42).

Normalized data had differential expression analysis undertaken with DESeq2 with comparisons across stages (simple models). To account for potential confounding by tissue composition, stromal and immune scores calculated using ESTIMATE were also included as covariates (adjusted models). Differential expression trends observed across increasing disease severity were highly consistent between the simple and adjusted models. Significant correlation (p < 2.2e-16) was found between log2 fold changes across the simple and adjusted models (Supplementary Figure S1D-G). Further, over 75% of up and downregulated genes overlapped across the two models for all comparisons, indicating similarity in overall direction and magnitude (Supplementary Table S3). These findings suggest that the tissue composition of the samples had minimal influence on differential expression analysis findings. Therefore, we have used differential expression analysis results from the simple model for examining the global transcriptomic profiling trends and pathway analysis. For biomarker discovery, we opted to be more conservative by focusing on genes that are significantly differentially expressed in both models.

### Global transcriptomic alterations reveal early-divergence and progressive tissue heterogeneity

We performed a principal component analysis (PCA) using all expressed genes for multivariate analysis and visualization across samples from different stages. Across PC1, NSq samples clustered separate from disease stages, whereas along PC2, a gradual separation of NDBE, dysplasia (LGD/HGD), and EAC samples was observed (Figure 1A). Further multivariate analysis was undertaken with correlation heatmaps, generated based on the normalized expression values of the top 500 most variable genes. This orthogonal hierarchical clustering-based method further confirmed this trend. NSq samples exhibited a high similarity to one another and were clearly distinct from the disease stage samples. Additionally, the heatmap revealed shared gene expression patterns among disease stages, particularly for NDBE, LGD, and HGD (Figure 1B).

**Figure 1:**
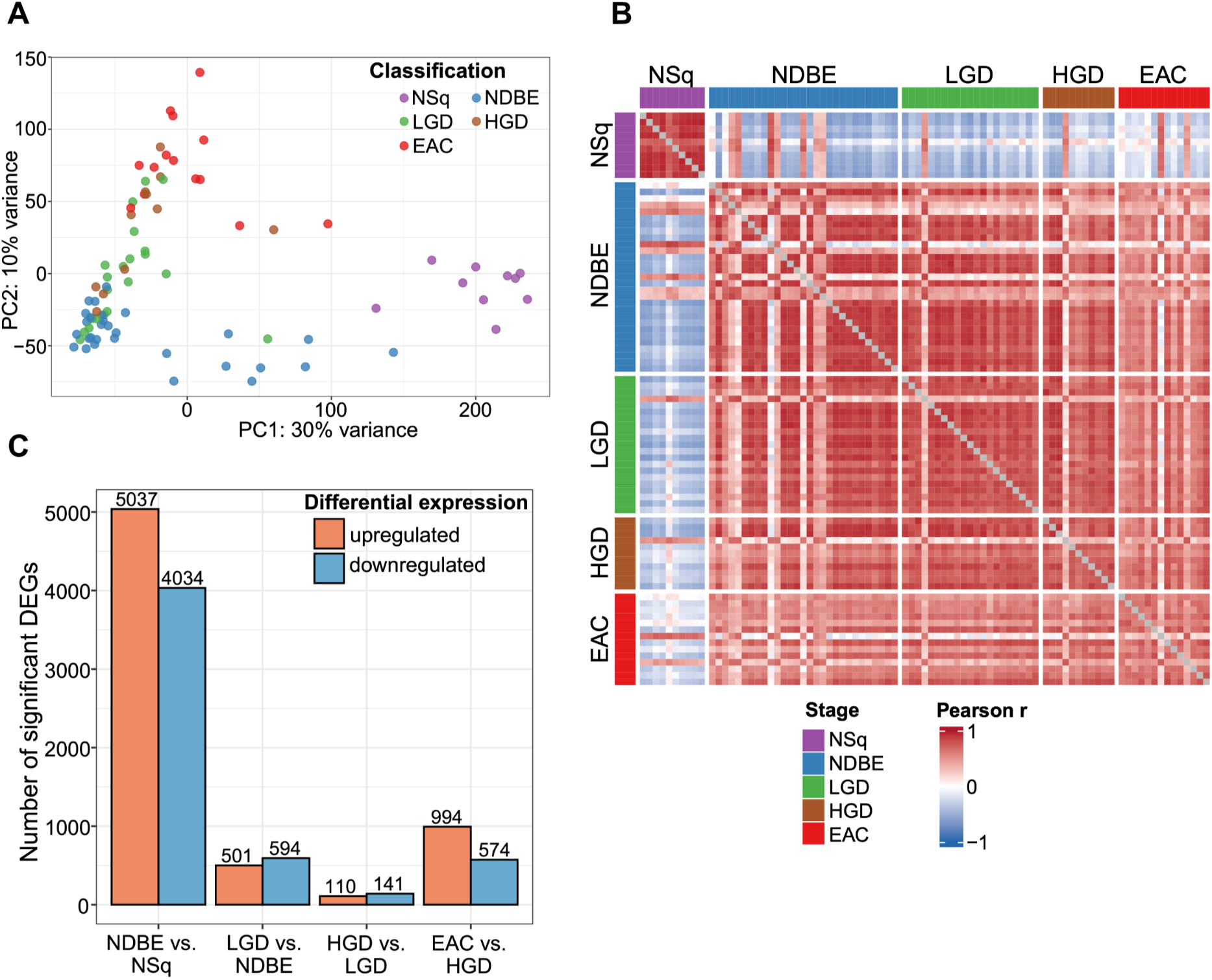
Overview of differential expression analysis. **(A)** Principal component analysis (PCA) of the samples using all expressed genes. First two components (PC1, PC2) are shown. **(B)** Pairwise Pearson correlation heatmap based on DESeq2-normalized expression values of the top 500 most variable genes. Samples are grouped by disease classification. **(C)** Bar plot showing the number of significantly upregulated (orange) and downregulated (blue) differentially expressed genes (absolute value(log2FC) > 1, p.adj < 0.05) identified for each pair-wise comparison across worsening disease stages using DESeq2. Colors and abbreviations are standardized across subplots. **NSq:** normal squamous, purple; **NDBE:** Non-dysplastic Barrett’s esophagus, blue; **LGD:** low-grade dysplasia, green; **HGD:** high-grade dysplasia, brown; **EAC:** esophageal adenocarcinoma, red.

A similar trend was observed in the number of significantly differentially expressed genes (DEGs) across disease progression stages (Figure 1C). The highest number of DEGs was detected in the pairwise comparison between NDBE and NSq (N=9071; |log2 fold change| > 1, adjusted p < 0.05). The number of DEGs was lower in subsequent comparisons, including LGD vs NDBE (N=1095), reaching the lowest in HGD vs LGD (N=251). A slight increase was observed for EAC vs HGD comparison (N=1568).

### Cell type transitions, increased proliferation in dysplasia and immune system dysregulation in EAC

We performed GSEA using GO terms alongside activity inference using PROGENy model (36) and CollecTRI resource (37) via decoupleR to gain complementary insights into transcriptional and signaling alterations across increasing disease severity (Figure 2A-C). While GSEA identified coordinated changes in gene expression patterns within predefined biological processes (33), decoupleR analysis enabled the inference of pathway and TF perturbations by evaluating experimentally derived downstream transcriptional responses, taking post-translational modifications into consideration (35).

**Figure 2:**
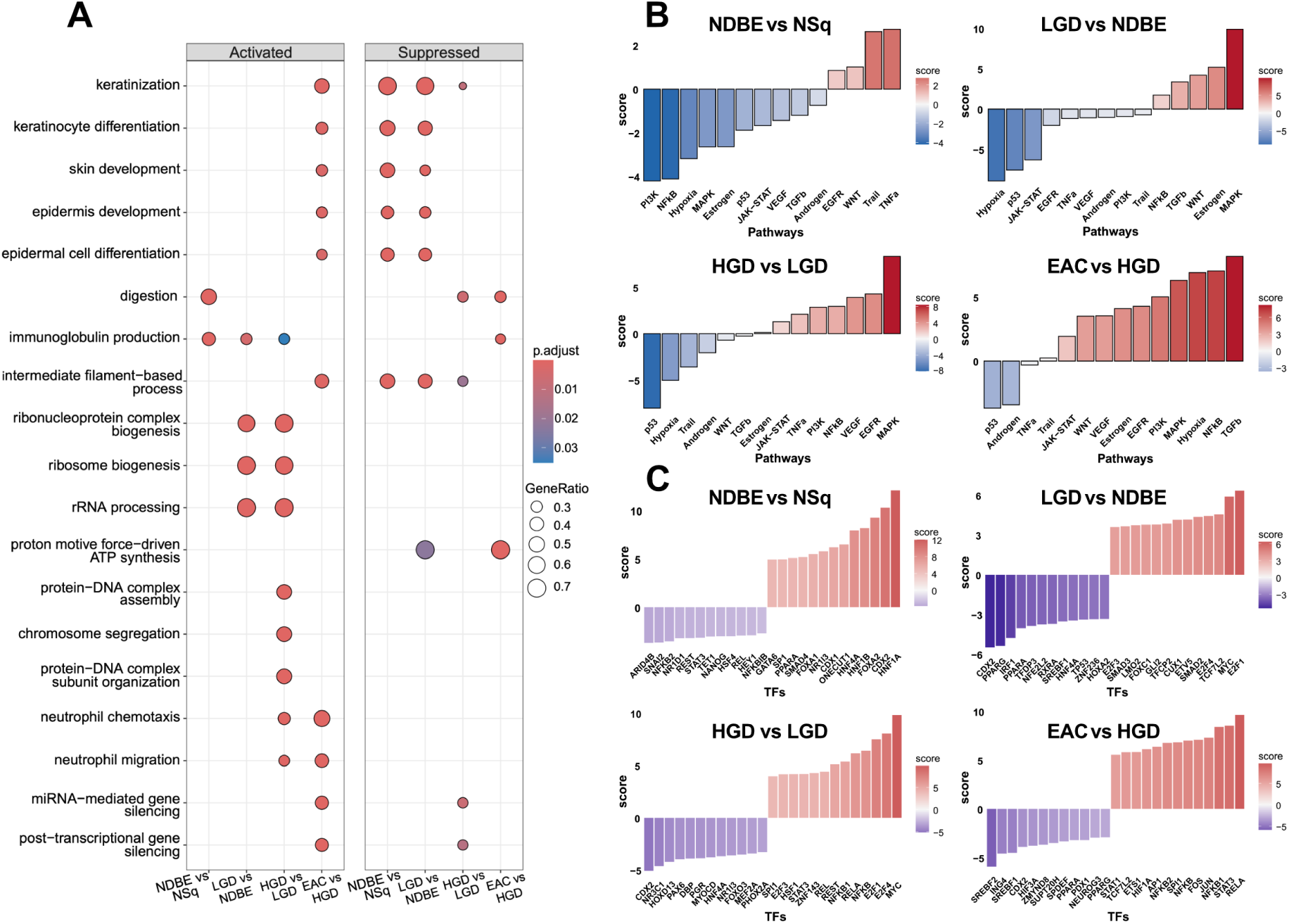
Functional enrichment analyses across increasing disease severity. **(A)** Gene Set Enrichment Analysis (GSEA) of Gene Ontology (Biological Process) gene sets across pairwise disease comparisons. Top significantly enriched pathways, both activated and suppressed, are shown. Dot size corresponds to the number of genes found to be significantly differentially expressed compared to the total number of genes in the specific pathway. Dot color represents the statistical significance. **(B)** Multivariate Linear Model (MLM) activity inference of 14 core signaling pathways in Pathway RespOnsive GENes (PROGENy) model based on downstream gene expression changes, for each pair-wise comparison. Bars represent the calculated t-statistic for each pathway. Positive scores denote pathway activation while negative scores indicate pathway repression. **(C)** Univariate Linear Model (ULM) activity inference of top 25 significantly different transcription factors (TFs) based on the expression of their downstream target genes annotated in the CollecTRI resource, for each pairwise comparison. Bars represent the calculated t-statistic for each TF. Positive scores denote TF activation while negative scores indicate TF repression.

During the metaplastic transition from NSq to NDBE, GO biological processes associated with epidermis development and keratinocyte differentiation, which are characteristic of the NSq epithelium, were significantly suppressed, while digestion-related processes were activated (Figure 2A). A similar trend was observed in over- representation analysis (ORA) performed on up and downregulated genes common for pairwise comparisons of disease stages (NDBE, LGD, HGD, EAC) with NSq (Supplementary Figure S2). These findings support the current notion of distinct cellular origin in BE (22) and suggest a shared gene expression program associated with cell type difference, specifically the transition from squamous epithelium to columnar cells characteristic of intestinal metaplasia. This was accompanied by a notable predicted increase in homeobox transcription factor CDX2 activity, further supporting the establishment of intestinal lineage identity (Figure 2C and Supplementary Figure S3) (22,43). PROGENy analysis revealed an increase in the apoptotic pathway TRAIL (Tumor necrosis factor (TNF)-related apoptosis-inducing ligand) and TNF-α (Tumor necrosis factor-alpha), along with a reduction in NF-kB signaling and proliferative and stress-associated signaling, including MAPK/PI3K pathways (44) (Figure 2B).

The transition from NDBE to LGD to HGD was associated with notable activation of cell proliferation associated biosynthetic and mitotic processes, including ribosome biogenesis, rRNA processing, chromosome segregation and protein-DNA complex organization. This transition was further associated with increased activity of transcription factors MYC and TFCP2 consistent with increased cellular growth and division during neoplastic progression (45). CDX2 activity declined in dysplasia compared to non-dysplastic Barrett’s further highlighting the progression from intestinal metaplasia to a more proliferative precancerous state.

Comparisons between stages from NDBE to dysplasia and EAC were marked by upregulation of immune system processes including neutrophil chemotaxis and migration, further supported by an increase in NF-kB signaling pathway and upregulation of its transcription factor components NFKB1, NFKB2 and RELA. Notably, during the transition from HGD to EAC, immunoglobulin production was reduced, while neutrophil chemotaxis and migration further increased, suggesting the development of an immunosuppressive environment (46). Hypoxia signaling declined across NDBE to dysplasia followed by a sharp increase in EAC samples. Oxygen deprivation and hypoxia are known to correlate with immune cell infiltration in EAC (47), and fits with the higher immune cell presence observed in this study.

Moreover, we observed a coordinated activation of developmental and oncogenic signaling pathways including Wnt, TGF-b, MAPK, PI3K and VEGF (44). Progressive disruption of p53 pathway was observed from NDBE to dysplasia to EAC, consistent with TP53 mutations, loss of heterozygosity and functional inactivation reported in previous studies (45,48). Furthermore, HGD to EAC transition was associated with reversal of epithelial differentiation and digestive programs that were established during NSq to NDBE transition.

### Transcriptomic profiling identifies candidate biomarkers for distinction of dysplasia and EAC

Based on differential expression analysis across disease stages, we aimed to identify a list of potential biomarkers that could distinguish tissues with dysplasia and adenocarcinoma from NDBE at the transcript level. The intent was to find biomarkers that assist with the often-difficult histopathological diagnosis of dysplasia, and that help identify patients for whom Barrett’s eradication therapy may be beneficial.

We identified a total of 574 genes that were significantly differentially expressed in dysplasia/EAC compared to NDBE. This group consisted of 110 upregulated (97 protein-coding, 13 ncRNA) and 464 downregulated (185 protein-coding, 279 ncRNA) genes (Supplementary Table S4). Multivariate analysis was then used to examine both the diagnostic utility of this subset and the correlation of these markers across disease stages. Figure 3A shows hierarchical unsupervised clustering of 85 samples from the discovery cohort based on log2-normalized expression of the selected biomarkers. Despite some inconsistencies, the samples largely clustered together according to disease stage.

**Figure 3:**
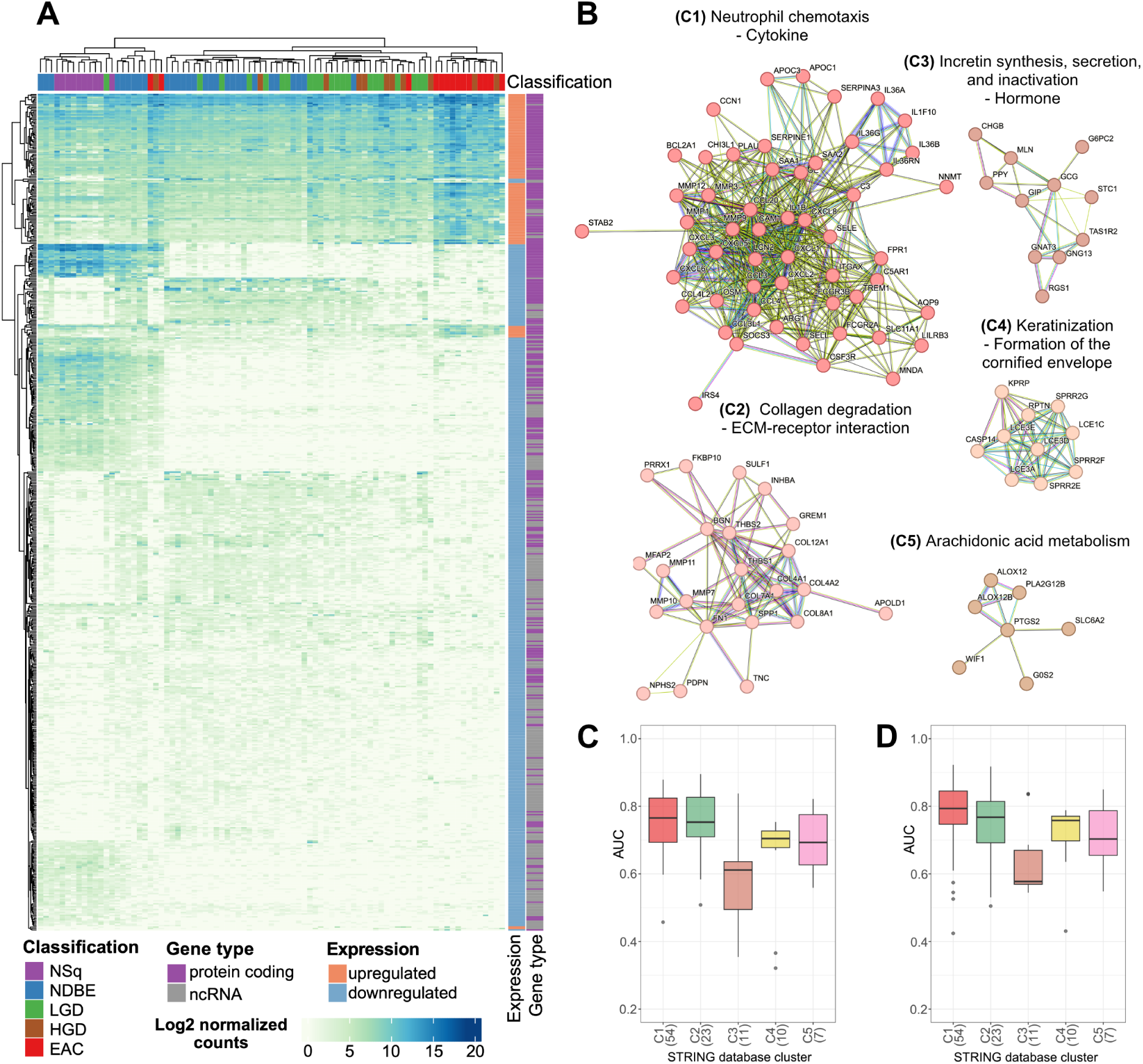
Network-based identification of hub biomarker candidates associated with dysplasia and esophageal adenocarcinoma (EAC). **(A)** Heatmap showing unsupervised hierarchical clustering of samples based on log2 normalized transcript counts of shortlisted biomarkers, comprising 110 upregulated and 464 downregulated genes associated with dysplasia and EAC. **(B)** Top five protein-protein interaction (PPI) clusters identified from the shortlisted biomarkers using the STRING database and Markov Cluster (MCL) algorithm clustering. Only protein-coding genes were included in the network analysis. **(C-D)** Receiver operating characteristic (ROC) analysis showing the area under the curve (AUC) values for individual protein coding genes within the top five STRING PPI clusters. Number of genes within each cluster is indicated within brackets. **(C)** Discovery cohort (n=85) **(D)** Independent validation cohort (published data from Maag et al., 2017, n=51). The corresponding median AUC values for the discovery and validation cohorts are as follows; C1 (AUC=0.77 and 0.79), C2 (AUC=0.75 and 0.77), C3 (AUC=0.61 and 0.58), C4 (AUC=0.70 and 0.76) and C5 AUC=0.69 and 0.70).

Protein-Protein interaction (PPI) network functional enrichment analysis was then used as a further tool to find mechanistically linked biomarkers. Several of these protein-coding gene biomarker candidates emerged as hub proteins in STRING network analysis (38), highlighting their biological relevance (i.e. they are functionally linked, rather than random) and their potential role in disease progression to dysplasia and EAC (Figure 3B). The top 5 clusters identified in PPI network analysis using the Markov Cluster Algorithm with default input options (inflation parameter = 3) were enriched for neutrophil chemotaxis (cytokine, C1), collagen degradation (ECM- receptor interaction, C2), incretin synthesis, secretion and inactivation (hormone, C3), keratinization (formation of the cornified envelope, C4) and arachidonic acid metabolism (C5) (Supplementary Table S5).

To understand the significance of these clusters in the diagnosis of disease, the area under the receiver operating characteristic (ROC) curve (AUC) values, reflecting the ability of genes within these clusters to distinguish dysplasia and EAC from NSq/NDBE, were calculated for both discovery and validation (24) cohorts (visualized in the boxplots in Figure 3C-D).

### Machine learning prioritization refines a candidate biomarker panel

From the shortlisted candidates described above, we performed machine learning (ML)-based feature ranking to identify key biomarkers based on their ability to distinguish between NSq/NDBE (control) and dysplasia/EAC (case) groups. Variable selection was performed using both LASSO regression and XGBoost models, the top 20 features from each method were extracted, and a consensus list generated based on combined ranks. The diagnostic performance of the selected biomarkers was assessed by calculating AUC in the discovery and validation cohorts separately (Supplementary Tables S6 and S7).

The diagnosis of LGD is limited by substantial pathologist interobserver variability and LGD is frequently over-diagnosed in the community setting (49). Given this diagnostic uncertainty, the analysis was performed twice by alternately including LGD samples within the control and case categories. Both ML ranking runs yielded 32 genes each, with 8 genes in common between the two runs, resulting in a total of 56 unique genes (Supplementary Tables S6 and S7). Among these, 44 were protein-coding, with 30 of these present within the top 5 clusters generated in STRING database analysis. These were subsequently prioritized for IHC validation studies. Given that most ncRNA do not produce a protein product, the 12 non-coding genes were not pursued further for IHC analysis.

### Selection of a minimal biomarker panel for RNAseq based diagnostic applications

While IHC is standard practice in molecular pathology, targeted RNA-seq of FFPE samples is emerging as a diagnostic paradigm (50). Thus, we aimed to identify a suitable panel of genes for a potential RNAseq-based diagnostic test. The panel optimization was performed by starting with the top-ranked marker for NSq/NDBE vs LGD/HGD/EAC comparison (Figure 4A, Supplementary Table S6), and iteratively adding the next feature that maximizes the resulting logistic model cross-validated AUC until the improvement falls below a certain threshold. For this use case, there was no need to first filter out ncRNA genes or enrich for upregulated genes. The iterative inclusion of additional markers resulted in a five-gene panel including 2 protein-coding (SLC11A1, IL36A) and 3 non-coding RNA (lncRNA LUCAT1, miRNA MIR215, snRNA RNU6-954P) genes.

**Figure 4:**
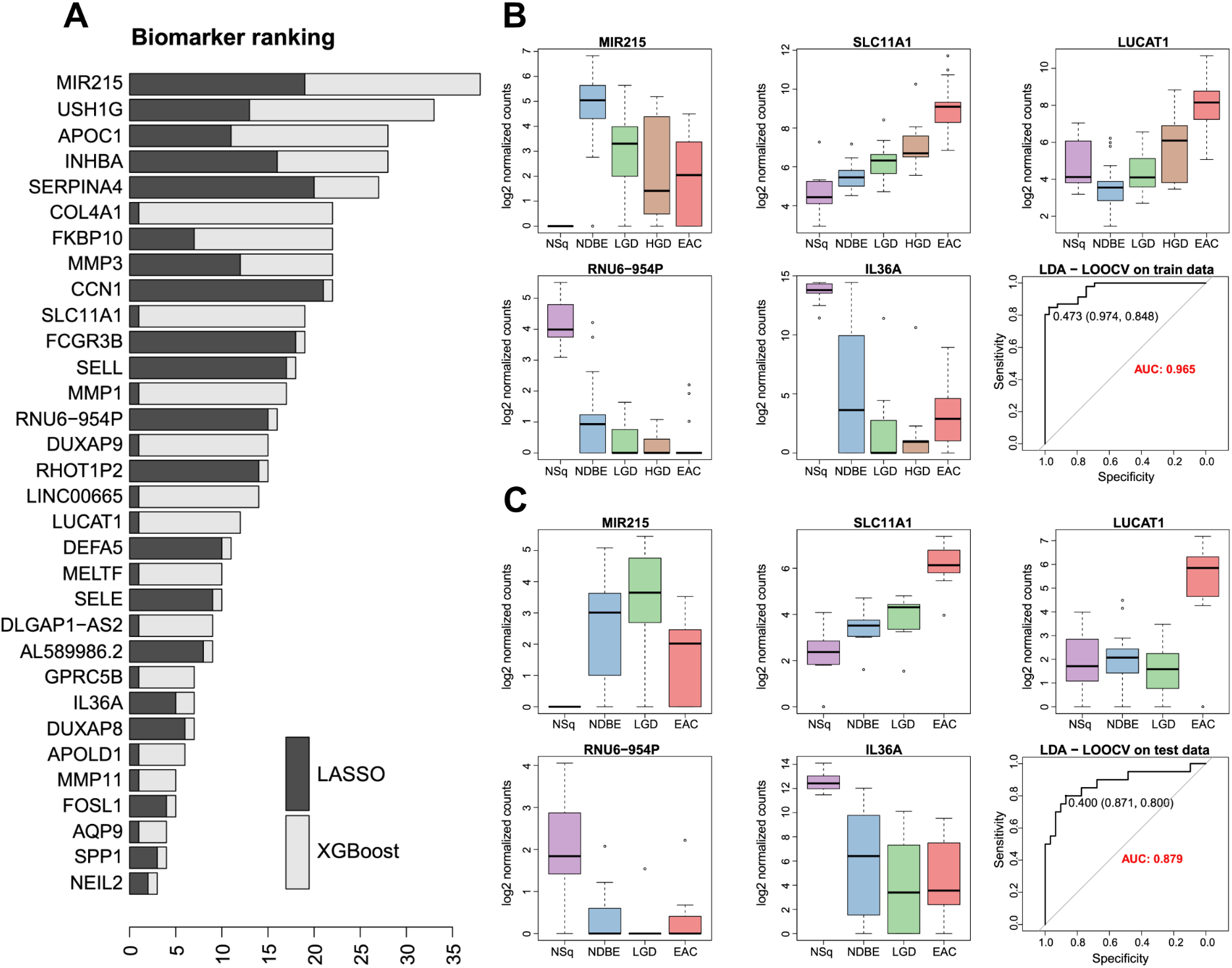
Selection of a gene panel for potential RNA sequencing-based diagnostic applications. **(A)** Combined gene ranking generated using LASSO and XGBoost for distinguishing LGD/HGD/EAC (case) from NSq/NDBE (control). **(B-C)** Diagnostic performance of the selected five-gene panel, comprising two protein-coding (SLC11A1, IL36A) and three non-coding RNA (long non-coding RNA LUCAT1, microRNA MIR215, small nuclear RNA RNU6-954P) genes **(B)** Receiver operating characteristic (ROC) analysis in the discovery cohort (training set) **(C)** ROC analysis in the independent validation cohort (test set). Performance is reported as the area under the curve (AUC), demonstrating the discriminative ability of the panel for classifying control and case samples. Colors and abbreviations are standardized across subplots. **NSq:** normal squamous, purple; **NDBE:** Non-dysplastic Barrett’s esophagus, blue; **LGD:** low-grade dysplasia, green; **HGD:** high-grade dysplasia, brown; **EAC:** esophageal adenocarcinoma, red

In the discovery cohort, the panel demonstrated strong classification performance with an AUC of 0.97, 97% specificity, and 85% sensitivity. The panel outperformed individual markers, which showed AUCs ranging from 0.67 to 0.88 (Figure 4B). The panel was evaluated in the independent validation cohort, where it achieved an AUC of 0.88 and a specificity of 87% and a sensitivity of 80% (Figure 4C). These results support the use of a multi-marker diagnostic panel over a single gene for improved performance.

### Protein biomarker expression in formalin-fixed paraffin-embedded (FFPE) tissue

From the list of candidates selected through ML-based ranking, seven protein-coding genes were chosen for IHC validation (Supplementary Table S6 and S7). These experiments aimed to assess whether directional expression patterns of the selected markers were present at the protein level in FFPE tissue and to technically validate suitable antibodies for IHC to take forward to a comprehensive future study with BE tissues; the present work was limited to tissue microarrays (TMAs) containing only NSq and EAC tissues. The selection criteria further included: (1) minimal expression in the control group (NSq/NDBE) to ensure low background staining (assessed by violin plots of log2 expression values in both discovery and validation cohorts, Supplementary Figure S4), (2) availability of antibodies validated for human tissue, and (3) prior reporting in gastrointestinal IHC studies. Due to the lack of validated monoclonal antibodies in human FFPE tissue, proteins SLC11A1 and IL36A from the above selected panel were excluded from IHC validation.

The selected IHC targets consisted of matrix metalloproteinases (MMP1 and MMP3), chemokines (CXCL5 and CCL3), cytokine oncostatin M (OSM), transforming growth factor-β superfamily protein inhibin (INHBA), and the cell surface receptor protein

Triggering Receptor Expressed on Myeloid cells 1 (TREM1). In addition, the downregulated biomarker Motilin (MLN) was included as it has the property of being highly expressed in Barrett’s sections at RNA level, but with a gradual reduction across the Barrett’s to neoplasia spectrum and very limited expression in many EACs. The biomarker PTGS2, which encodes for the protein cyclooxygenase-2 (COX-2), was also included as a well characterized BE/EAC marker and part of a commercially available diagnostic panel (51). Both MLN and PTGS2 are also present in top STRING clusters. Alongside these, we also assessed three routinely used clinical markers, tumor suppressor p53 (TP53), epidermal growth factor receptor 2 (HER2/ERBB2), and proliferation marker Ki-67 (MKI67).

The IHC results indicated that TREM1 (median of malignant vs normal; 45 vs 0, Wilcoxon rank sum test with continuity correction p < 0.0001), OSM (4 vs 0, p < 0.0001), and COX-2 (80 vs 0, p < 0.0001) were significantly overexpressed in malignant tissue compared to NSq. For CXCL5, although median H-score was 0 in both groups, the distribution of H-scores differed, with higher values in EAC tissues (p < 0.01, Wilcoxon rank sum test with continuity correction). In contrast, MLN was significantly downregulated in EAC relative to NSq (40 vs 240, p < 0.0001). Based on aberrantly high or low staining intensity (ROC-derived cutoff) in 56 malignant samples, TREM1 correctly identified 36 (64%), OSM identified 30 (54%), COX-2 identified 38 (68%), CXCL5 identified 17 (30%), and MLN identified 55 (98%). The expression of MMP1 and MMP3 showed no significant difference between two tissue types at the protein level (Supplementary Figure S5). INHBA and CCL3 staining were negative in both tissue types.

Aberrant p53 staining was present in 49/56 cases (88%), with 33 showing strong positive nuclear staining and 16 showing complete absence of p53 expression (Supplementary Figure S6). Ki-67 demonstrated significant overexpression in malignant tissue compared to NSq (median 26.5 vs 0, p < 0.0001), while correctly identifying 49 (88%) samples. HER2 was only positive in five (9%) malignant samples. Of the seven malignant cases that exhibited wild type p53 staining patterns (i.e. neither strongly positive nor completely absent), the novel markers were able to correctly identify these as follows; Motilin 7/7, TREM1 5/7, OSM 4/7, and COX-2 4/7. Ki-67 also showed positive staining in 5/7 cases. Notably, the two EAC cases missed by both p53 and Ki-67 were detected by OSM, TREM1 and MLN.

## Discussion

In this study, we performed comprehensive transcriptomic profiling across the full spectrum of Barrett’s disease. By evaluating stage-specific differences, we delineated gene expression alterations that accompany disease progression and identified candidate biomarkers with potential utility for distinguishing LGD, HGD, and EAC from NDBE and NSq.

Our findings demonstrate that the most extensive transcriptional changes occur during the transition from NSq to NDBE, marked by loss of squamous epithelial differentiation program alongside activation of gastrointestinal digestive functions. These results align with previous reports and highlight the cell type transformation during metaplastic progression (15,22). Compared to NDBE, LGD and HGD were characterized by enhanced cellular growth and proliferation programs, consistent with advancing neoplastic progression (45). These changes were further accompanied by upregulation of immune system processes and oncogenic signaling pathways during the transition from dysplasia to EAC (44,46). We also observed that HGD to EAC transition was associated with reversal of epithelial differentiation and digestive programs that were established during NSq to NDBE transition. This shift from HGD to EAC, likely reflects progressive dedifferentiation and changes in transcriptional programs from lineage specific metaplasia to malignant transformation (17).

Biomarker discovery further confirmed the dysregulation of biological processes associated with the increasing disease severity from NDBE to dysplasia and EAC. The two largest enriched biomarker clusters were neutrophil chemotaxis (cytokine) and collagen degradation (ECM-receptor interaction). These pathways reflect key features of Barrett’s carcinogenesis, with cytokine signaling driving immune cell recruitment (52), and extracellular matrix remodeling promoting epithelial plasticity, invasion and malignant transformation (53). The genes in these two clusters demonstrated notable discriminatory power in both discovery and validation cohorts, distinguishing dysplasia and EAC cases from NDBE/NSq, with median AUC values greater than or equal to 0.75.

Given the recent improvements in sequencing technologies, RNA-seq and targeted panel sequencing have emerged as attractive diagnostic approaches (50), complementing routine histopathology (54). Targeted panel sequencing could help identify dysplasia or EAC that was missed or uncertain by histopathological review, or to flag tissues for expert review. To date, there are no RNA-based tissue tests for the diagnosis of Barrett’s associated dysplasia and EAC. The most comprehensive RNA test currently available is the EMERALD test, which is based on circulating microRNA in blood (55). Previous studies, including our own (17,24), have identified potential tissue-based RNA biomarkers for diagnosis, but these studies have primarily focused on the comparison between EAC and NDBE.

In this study, we used machine learning to identify a five-gene panel comprising both protein-coding and non-coding RNAs for distinguishing dysplasia and EAC from NDBE and normal squamous epithelium in biopsy samples. Importantly, this panel was developed by considering gene expression patterns across the entire disease spectrum rather than between individual disease stages. The five-gene panel included 2 protein-coding (SLC11A1, IL36A) and 3 non-coding RNA (lncRNA LUCAT1, miRNA MIR215, snRNA RNU6-954P) genes.

MicroRNA (miRNA) MIR215 is well characterized in BE and EAC. MIR215 expression is consistently reported to be upregulated in BE compared to normal squamous esophageal tissue. In most studies, MIR215 expression gradually decreases relative to NDBE tissue during progression to worse BE stages, suggesting an association with IM and potential as a biomarker for early BE (56,57).

SLC11A1 (solute carrier family 11 member 1) gene encodes iron transporter protein Natural Resistance-Associated Macrophage Protein 1 (NRAMP1) which is primarily expressed in macrophages. NRAMP1 is known to play a role in modulating tumor microenvironment and immune response and elevated levels have been implicated in a variety of cancers including esophageal squamous cell carcinoma (ESCC) (58). Interleukin-36 alpha (IL-36α) encoded by IL36A gene is an inflammatory cytokine that is abundantly expressed in healthy esophageal epithelial tissue (59). IL-36α appears to function as a tumor suppressor in several cancer types, including hepatocellular, ovarian and colorectal, with higher expression associated with improved survival while reduced expression correlated with disease progression and poor prognosis (60).

LUCAT1 (lung cancer associated transcript 1) is an oncogenic lncRNA that is frequently upregulated in a range of cancers, including ESCC. Given the observed correlation between its expression levels and clinical parameters such as TNM staging, histopathology grade, tumor size and overall survival, LUCAT1 has emerged as a potential diagnostic and/or prognostic biomarker and a candidate therapeutic target (61). Small nuclear RNA (snRNA) RNU6-954P is poorly characterized in cancer biology.

While RNA-seq could be promising, H&E staining is the pathology clinical standard (62), and thus highly compatible with IHC ancillary tests. Therefore, we investigated the expression of IHC candidate markers at protein level in malignant (EAC and GEJ adenocarcinoma) and normal squamous tissues in commercially available tissue microarrays to determine whether the expression trends observed at transcript level correlated with protein expression.

The selected markers OSM, CXCL5, TREM1 and COX-2 exhibited protein expression patterns consistent with those observed at transcript level, with significantly higher expression in malignant tissue compared to NSq. A recent study of EAC and ESCC found that high OSM expression correlated with poor overall survival, advanced stage and lymph node metastasis (63). Pro-tumor chemokine CXCL5 promotes EAC progression by driving an immunosuppressive tumor microenvironment and angiogenesis (64). A recent pan-cancer study including EAC and ESCC, found that inflammatory marker TREM1 promoter hypomethylation leads to overexpression associated with poor survival (65). Although TREM1 has not been studied in BE specifically, it has been implicated in reflux esophagitis (66). Inflammatory enzyme COX-2 expression is progressively upregulated during the transition from BE to dysplasia to EAC (67), and it is incorporated in the commercially available TissueCypher test used for risk stratification (51). MLN expression was significantly higher in NSq compared to malignant tissue in contrast to the transcript level trend, suggesting motilin protein warrants for further evaluation in BE. Although previously not linked to BE/EAC, an indirect association may exist through motilin’s role in modulating gastric emptying and lower esophageal motility (68).

P53 and Ki-67 have demonstrated diagnostic utility in identifying presence of dysplasia, particularly in histologically challenging cases such as LGD and indefinite for dysplasia, along with predicting risk of progression (69,70). Ki-67 expression increases progressively along the Barrett’s metaplasia-dysplasia-adenocarcinoma sequence, while aberrant p53 staining in EAC reflects underlying TP53 mutation. Staining of both markers differed significantly between NSq and malignant tissue, with their combination resulting in only two cancers with staining that was not different from normal tissues. These two cases showed marked differences in TREM1, OSM and MLN staining. HER2, primarily used to identify candidates for targeted therapy in established EAC (71), was only positive in five malignant samples.

The inclusion of all Barrett’s stages for biomarker discovery using RNA-seq is a key strength of this study. However, the lack of prior studies following the same approach limited our ability to independently validate the proposed five-gene panel in external cohorts. Moreover, although we identified several IHC candidate markers and confirmed which maintained the expression trend at protein level, these markers were not assessed in BE samples. Consequently, their diagnostic utility in detecting different grades of dysplasia compared to NDBE remains to be established.

In summary, by performing transcriptome profiling across the full spectrum of Barrett’s disease, we characterized the molecular and biological processes underlying increasing disease severity, many of which are consistent with previous reports. We identified a five-gene panel signature that distinguishes dysplasia (LGD/HGD) and EAC from NDBE and NSq. We performed an initial validation study for this signature using the data from our previous RNA-seq study, but we acknowledge that further extensive validation is required before considering this panel for clinical use. In addition, we identified several potential IHC markers; future studies could include further evaluation of these, especially in Barrett’s tissues, and further validation of the RNA signature.

## Acknowledgements

The authors gratefully acknowledge the patients who provided samples for this study. This work was supported by National Health and Medical Research Council (NHMRC, Ideas Grant 2012513), St. Vincent’s Clinic Foundation and JW&M Cunnigham Foundation grant funding. We also thank Australian Genome Research Facility (Melbourne, Australia) for library creation and sequencing.

## Author contributions

J.R. and R.V.L conceived and designed the study. T.P.U, K.M.L. and R.V.L performed sample selection. T.Y., P.K., D.M. and I.B. conducted the pathology assessments. T.P.U., Y.J.L and K.M.L. carried out RNA extractions and quality control. T.Y. and G.G. performed the immunohistochemistry assays and/or scoring. T.P.U., Y.J.L., D.P., J.R. and R.V.L. carried out data analysis and/or interpretation. T.P.U. drafted the initial manuscript. T.P.U., Y.J.L., D.P., S.J.L., J.R. and R.V.L. reviewed and edited the manuscript. All authors read, provided feedback and approved the final version of the manuscript.

## Notes

**Conflict of interest statement**: The authors declare no potential conflicts of interest.

### Competing Interest Statement

The authors have declared no competing interest.

