## Supplementary Figures for "Transcriptomic Changes and Biomarkers in Barrett’s Metaplasia, Dysplasia and Cancer"

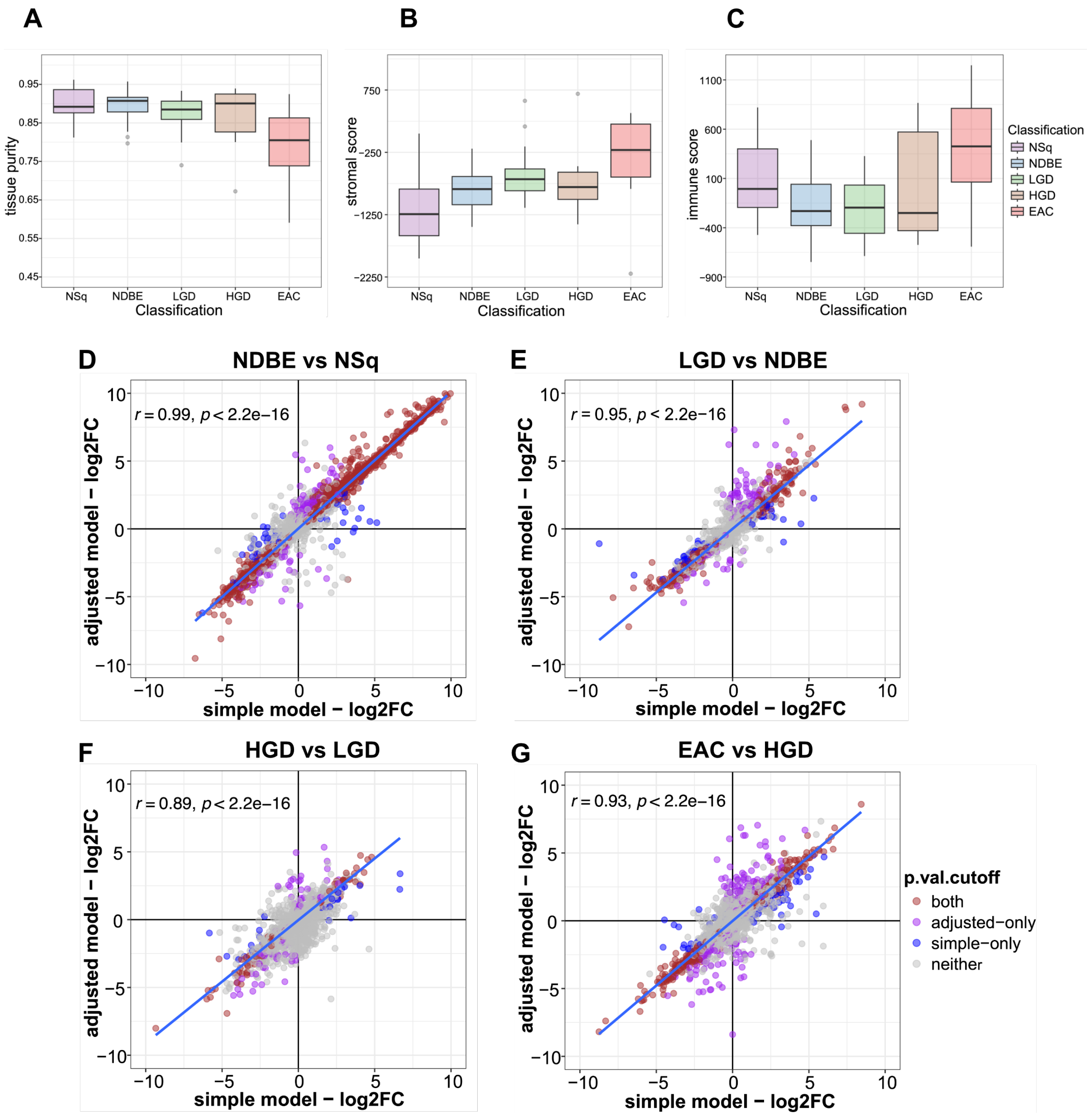


**Supplementary Figure S1: *In-silico* tissue purity estimations and comparison of differential expression models.**

Tissue purity was estimated using ESTIMATE (Estimation of STromal and Immune cells in MAlignant Tumor tissues) alogorithm. **(A)** Stromal scores **(B)** Immune scores **(C)** Estimated tissue purity for each sample group. **(D-G)** Correlation plots comparing log2 fold change values obtained from simple and adjusted models in DESeq2 for the pairwise comparisons **(D)** NDBE vs NSq **(E)** LGD vs NDBE **(F)** HGD vs LGD **(G)** EAC vs HGD. Dot color represents the significance of associated p-value (cutoff $p < 0.05$). Pearson's correlation coefficient is indicated for each comparison.

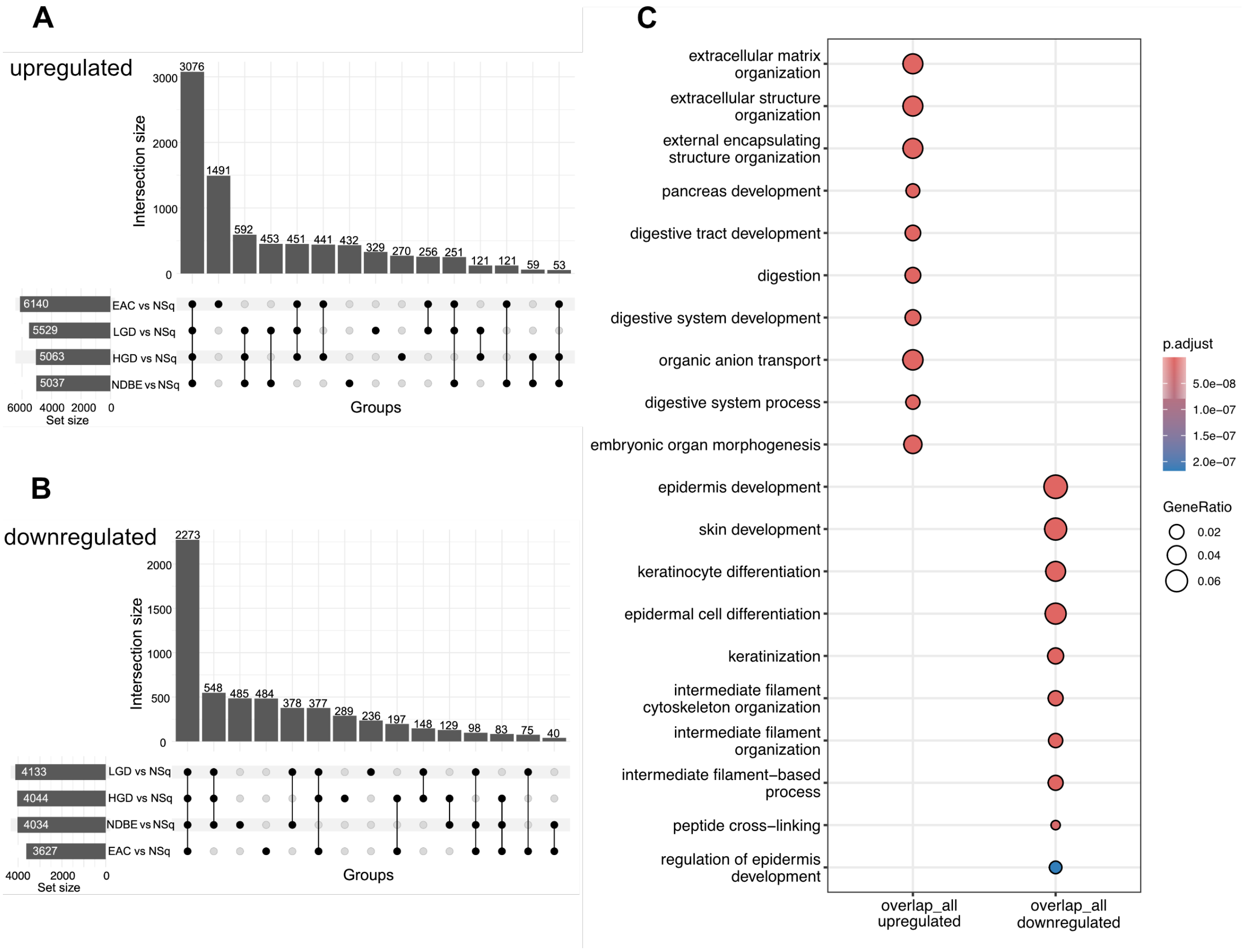


**Supplementary Figure S2: Differentially expressed genes shared among pairwise comparisons of disease stages with normal squamous (NSq) epithelium**

**(A-B)** UpSet plots showing the overlap of significantly **(A)** upregulated and (**B)** downregulated genes identified in pairwise comparisons of disease stages, non-dysplastic Barrett's esophagus (NDBE), low-grade dysplasia (LGD), high-grade dysplasia (HGD) and esophageal adenocarcinoma (EAC) with normal squamous (NSq). **(C)** Over representation analysis (ORA) of Gene Ontology (Biological Process) of the genes identified as differentially expressed (absolute value(log2FC)>1, p.adj<0.05) and shared by all disease stages compared to NSq. Top significant pathways, both activated and suppressed, are shown. Dot size corresponds to the number of genes found to be significantly differentially expressed compared to the total number of genes in the specific pathway. Dot color represents the statistical significance.

**A**

| Pathway | NDBE vs NSq | LGD vs NDBE | HGD vs LGD | EAC vs HGD |
|---|---|---|---|---|
| MAPK | ↓ | ↑ | ↑ | ↑ |
| Hypoxia | ↓ | ↓ | ↓ | ↑ |
| EGFR | NS | ↓ | ↑ | ↑ |
| p53 | NS | ↓ | ↓ | ↓ |
| NFkB | ↓ | NS | ↑ | ↑ |
| PI3K | ↓ | NS | ↑ | ↑ |
| Estrogen | ↓ | ↑ | NS | ↑ |
| Androgen | NS | NS | ↓ | ↓ |
| VEGF | NS | NS | ↑ | ↑ |
| TGFb | NS | ↑ | NS | ↑ |
| WNT | NS | ↑ | NS | ↑ |
| TNFa | ↑ | NS | ↑ | NS |
| Trail | ↑ | NS | ↓ | NS |
| JAK-STAT | NS | ↓ | NS | NS |

**B**

| TF | NDBE vs NSq | LGD vs NDBE | HGD vs LGD | EAC vs HGD |
|---|---|---|---|---|
| CDX2 | ↑ | ↓ | ↓ | ↓ |
| NFKB2 | ↓ | NS | ↑ | ↑ |
| PPARA | ↑ | NS | ↓ | ↓ |
| HNF4A | ↓ | ↑ | NS | NS |
| HNF1A | ↑ | NS | ↓ | NS |
| NR1I3 | ↑ | NS | ↓ | NS |
| TAL1 | ↓ | NS | ↑ | NS |
| TFAP2C | ↓ | NS | ↑ | NS |
| PDX1 | ↑ | NS | NS | ↓ |
| STAT3 | ↓ | NS | NS | ↑ |
| MYC | NS | ↑ | ↑ | NS |
| SOX2 | NS | ↓ | ↑ | NS |
| TFCP2 | NS | ↑ | ↑ | NS |
| ING4 | NS | ↓ | NS | ↓ |
| SREBF1 | NS | ↓ | NS | ↓ |
| TCF7L2 | NS | ↑ | NS | ↑ |
| NFKB | NS | NS | ↑ | ↑ |
| NFKB1 | NS | NS | ↑ | ↑ |
| RELA | NS | NS | ↑ | ↑ |

**Supplementary Figure S3: Inference of signaling pathway and transcription factor (TF) activity across disease progression.**

Summary of the activity of **(A)** 14 core signaling pathways in Pathway RespOnsive GENes (PROGENy) model based on downstream gene expression changes, for each pair-wise comparison and **(B)** top significantly altered TFs based on the expression of their downstream target genes annotated in the CollecTRI resource, for each pair-wise comparison. Activation (red) and repression (blue) are indicated by the direction and color of the arrows in the table.

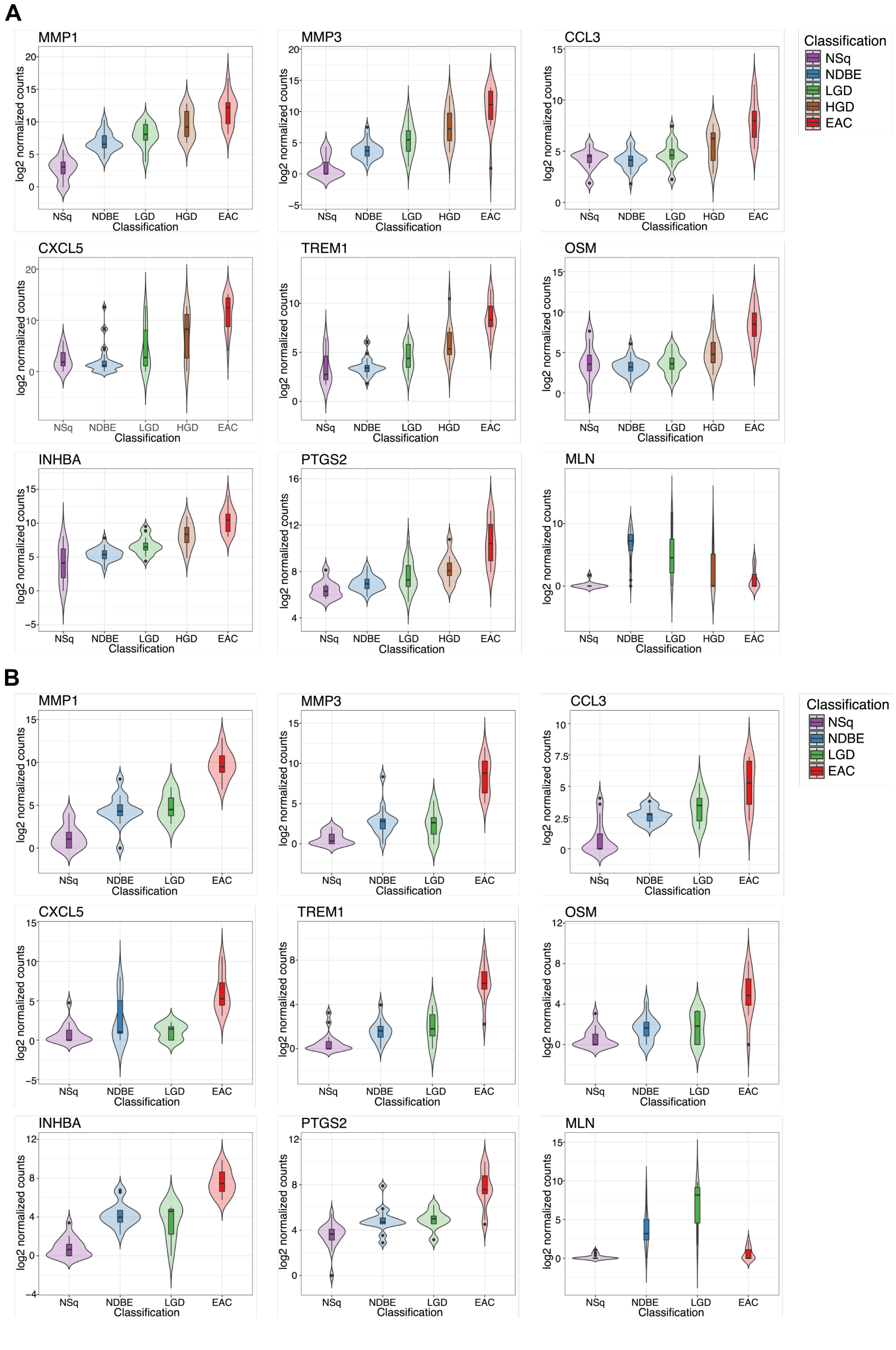


**Supplementary Figure S4: Expression of candidate immunohistochemistry (IHC) markers across disease stages.** Violin plots visualizing the normalized log2 transcript counts per each stage for the nine proteins selected as potential IHC targets. **(A)** Discovery cohort (n=85) **(B)** Validation cohort (published data from Maag et al., 2017, n=51) Colors and abbreviations are standardized across subplots. **NSq:** normal squamous, purple; **NDBE:** Non-dysplastic Barrett's esophagus, blue; **LGD:** low-grade dysplasia, green; **HGD:** high-grade dysplasia, brown; **EAC:** esophageal adenocarcinoma, red

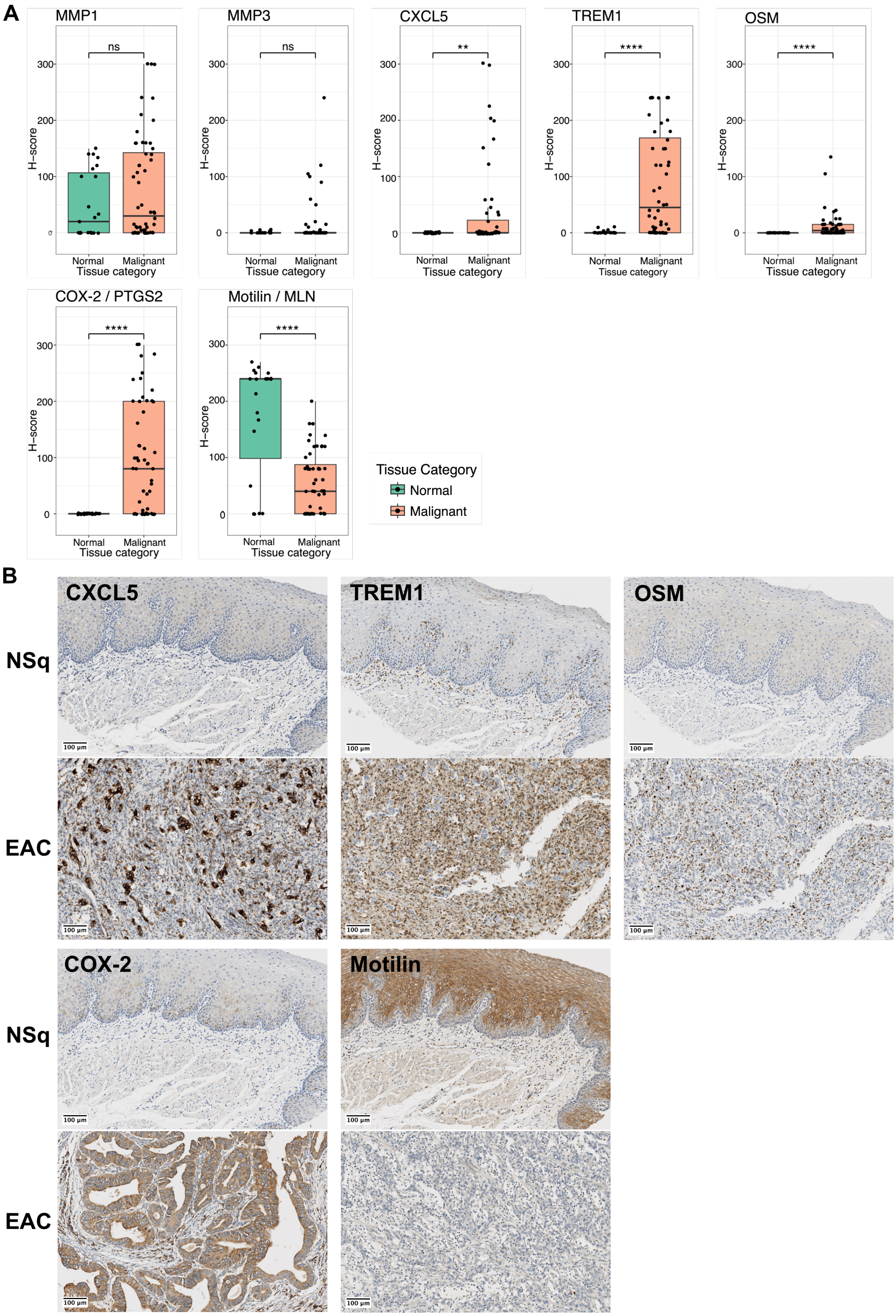


**Supplementary Figure S5: Immunohistochemical (IHC) validation of shortlisted biomarker proteins in formalin-fixed paraffin-embedded (FFPE) tissues. (A)** Box plots showing IHC staining scores for candidate biomarker proteins in normal and malignant tissue microarray (TMA) samples. Statistical significance was assessed using the Wilcoxon rank sum test with continuity correction. Significance levels are denoted as "****" ($p < 0.0001$), "***" ($p < 0.001$ ), "**" ($p < 0.01$), "*" ($p < 0.05$), and "ns" (not significant). **(B)** Representative IHC images of candidate biomarkers showing significantly differential protein expression between normal and malignant tissue. The top panel shows normal squamous (NSq) and the bottom panel shows esophageal adenocarcinoma (EAC) tissue samples.

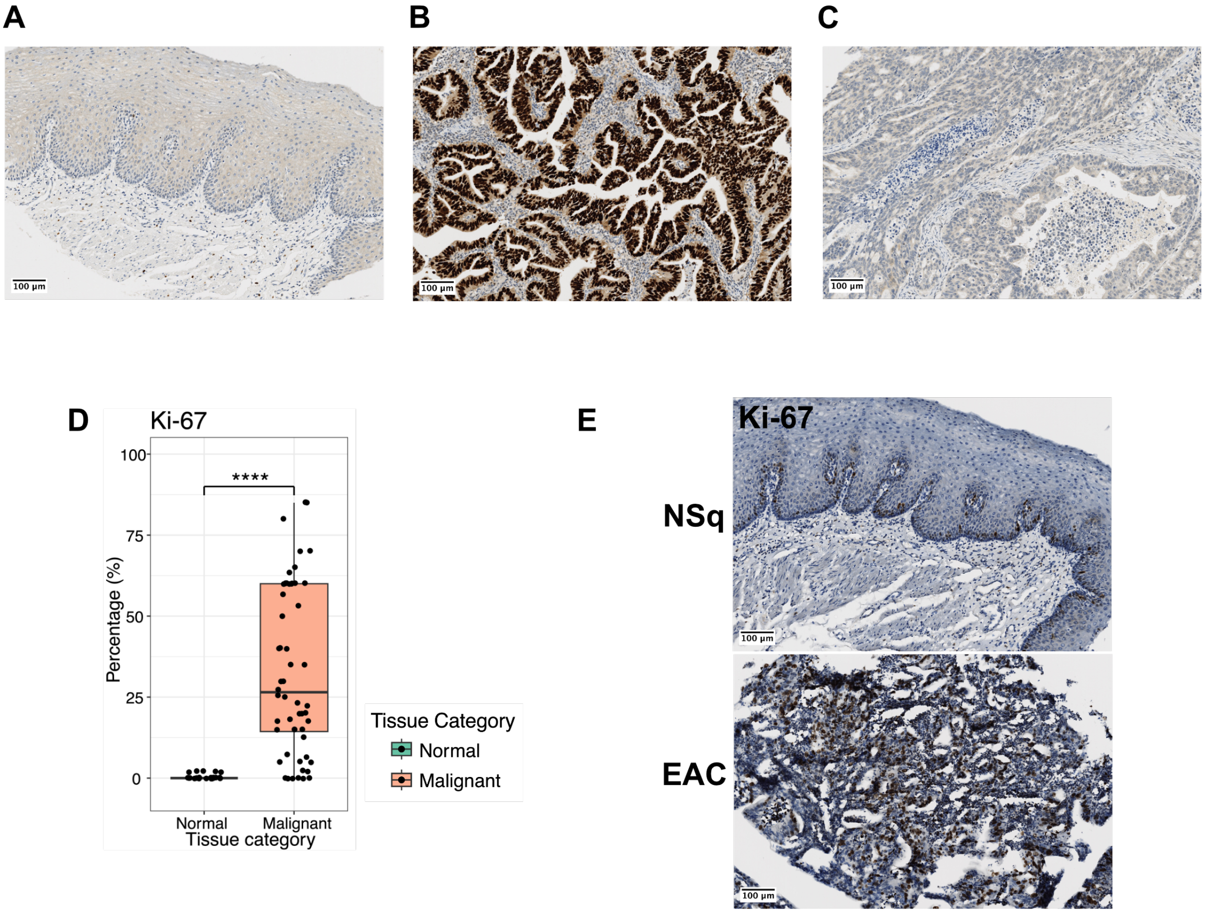


**Supplementary Figure S6: Immunohistochemical (IHC) analysis of clinical markers in formalin-fixed paraffin-embedded (FFPE) tissues.**

**(A-C)** Representative IHC images of p53 staining **(A)** Normal squamous (NSq) tissue **(B)** Esophageal adenocarcinoma (EAC) tissue with strong positive nuclear staining **(C)** EAC tissue showing complete absence of staining **(D)** Box plot showing Ki-67 staining indicated as percentage of cells in normal and malignant tissue microarray (TMA) samples. Statistical significance was assessed using the Wilcoxon rank sum test with continuity correction. Significance level "****" ($p < 0.0001$) **(E)** Representative IHC images of Ki-67. The top panel shows normal squamous (NSq) and the bottom panel shows esophageal adenocarcinoma (EAC) tissue samples.
